# D_1_- and D_2_-like receptors differentially mediate the effects of dopaminergic transmission on cost/benefit evaluation and motivation in monkeys

**DOI:** 10.1101/2020.11.27.400911

**Authors:** Yukiko Hori, Yuji Nagai, Koki Mimura, Tetsuya Suhara, Makoto Higuchi, Sebastien Bouret, Takafumi Minamimoto

**Affiliations:** Department of Functional Brain Imaging, National Institutes for Quantum and Radiological Science and Technology, Chiba, 263-8555, Japan; Team Motivation Brain & Behavior, Institut du Cerveau et de la Moelle épiniŕe (ICM), Centre National de la Recherche Scientifique (CNRS), Hôpital Pitié Salpêtriŕe, 75013 Paris, France

**Keywords:** dopamine, reward delay, workload, PET imaging, non-human primates, decision making

## Abstract

It has been widely accepted that dopamine (DA) plays a major role in motivation, yet the specific contribution of DA signaling at D1-like receptor (D1R) and D2-like receptor (D2R) to cost-benefit trade-off remains unclear. Here, by combining pharmacological manipulation of DA receptors (DARs) and positron emission tomography imaging, we assessed the relationship between the degree of D1R/D2R blockade and changes in benefit- and cost-based motivation for goal-directed behavior of macaque monkeys. We found that the degree of blockade of either D1R or D2R was associated with a reduction of the positive impact of reward amount and increasing delay-discounting. Workload-discounting was selectively increased by D2R antagonism. In addition, blocking both D1R and D2R had a synergistic effect on delay-discounting but an antagonist effect on workload-discounting. These results provide fundamental insight into the distinct mechanisms of DA action in the regulation of the cost and benefit-based motivation, which have important implications for motivational alterations in both neurological and psychiatric disorders.

## Introduction

In our daily lives, we routinely determine whether to engage or disengage in an action according to its benefits and costs: the expected value of benefits (i.e., rewards) has a positive influence, while the cost necessary to earn the expected reward (e.g., delay, risk or effort) decreases the impact of reward value [1–3]. Arguably, the dopamine (DA) system plays a central role in the motivation, which adjusts behavior as a function of expected costs and benefits. Phasic firing of mid-brain DA neurons positively scales with the magnitude of future rewards, and negatively scales with risk or time delay to reward [4–11]. In addition, several studies demonstrated that DA neurotransmission was causally involved in regulation of behavior based on expected costs and benefits [12–18]. In patients suffering from depression, schizophrenia or Parkinson’s disease (PD), the alteration of DA transmission is frequently associated with various pathological impairments of motivation such as anergia, fatigue, psychomotor retardation, and apathy [14, 19–21]. DA signaling is mediated at post-synaptic sites by two classes of DA receptors (DARs), the D1-like receptor (D1R) and the D2-like receptor (D2R), and both classes are thought to be involved in the regulation of motivation [22, 23].

However, the specific mechanisms through which DA contributes to motivation based on cost-benefit trade-off remain unclear. For example, in tasks where animals must exert a higher force to obtain a bigger reward, blockade of either D1R or D2R shifts preferences towards less efforts, thus less rewards, suggesting a role of DA in effort [24–29]. On the other hand, since DA activity shows little sensitivity to information about effort when it is decoupled from reward, it has been proposed that DA is strongly involved in adjusting motivation based on expected benefits (reward availability) rather than on expected energetic costs (effort) [9, 30, 31]. Note that this apparent controversy might be related to the difficulty of interpreting results from experiments where the nature of costs and benefits were not clearly identified and isolated [11].

To understand the role of DA in motivation, it is critical to identify not only the pattern of DA activity and release across costs and benefits, but also the action of DA on DARs [17]. However, the relative implication of distinct receptor sub-types in specific aspects of the cost-benefit trade-off in motivation also remains under debate. For example, systemic administration of D1R or D2R antagonist was shown to increase preference for small immediate rewards over larger, delayed rewards [25, 32–34]. Some of these studies, however, have also shown that blockade of D1R [34] or D2R [33] has no effect on delay cost. These and other previous behavioral pharmacology studies have compared the effect of DAR blockade according to the antagonist dose-response relationship for each DAR subtype. However, since different antagonists display distinct pharmacological properties (e.g., target affinity, brain permeability, biostability), it is difficult to accurately predict the effects on their target receptors in vivo. Therefore, to describe the role of DARs in motivational processes beyond a simple dose-response relationship, we considered it to be essential to measure receptor occupancy after antagonist administration. Indeed, positron emission tomography (PET) studies of patients have shown that in vivo D2R occupancy is a reliable predictor of clinical and side effects of antipsychotic drug s[35, 36]. Similarly, receptor occupancy has been measured in rats and monkeys, as well as the relationship with the behavioral effects following D2R antagonists [37–39].

In the present study, we aimed to quantify and directly compare the roles of DA signaling via D1R and D2R in motivation based on the costs and benefits in macaque monkeys. For this purpose, we manipulated DA transmission by systemic injections of specific antagonists for D1R and D2R and assessed the degree of DA receptor occupancy using in vivo PET imaging with selective radioligands. The effects of this quantitatively controlled DAR blockade on benefit- and cost-based motivation were evaluated in two sets of behavioral experiments. First, we quantified the effects of DAR blockade on the incentive impact of reward prediction, namely the relationship between predicted reward amount and the motivation of a goal-directed task. Second, to assess the effect of DAR blockade on two types of costs, workload and delay, we used a similar behavioral task for a fixed amount of reward, but either cost was implemented, allowing us to estimate the negative impacts of cost as steepness of reward discounting (i.e., workload- and delay-discounting). Based on our data, D1R and D2R have similar roles in incentive impact of reward prediction and delay discounting, whereas D2R is exclusively related to workload discounting.

## Results

### PET measurement of D1R/D2R occupancy following systemic antagonist administration

To establish appropriate antagonist doses and experimental timing, we measured the degree of receptor blockade (i.e., receptor occupancy) following systemic administration of DAR antagonists. We performed PET imaging with selective radioligands for D1R ([^11^C]SCH23390) and D2R ([^11^C]raclopride) in a total of 4 monkeys (3 for each) under awake condition for both baseline (without drug administration) and following antagonist administration. We quantified specific r adioligand b inding u sing a s implified reference tissue model with the cerebellum as reference region.

For D1R measurement, high radiotracer binding was seen in the striatum at baseline condition (Fig 1A, Baseline). PET scans were obtained after pre-treatment with non-radiolabeled SCH23390 for D1R antagonist at different doses (10, 30, 50, and 100 μg/kg), demonstrating that specific tracer binding was diminished in a dose-dependent manner (Fig 1A). We performed a volume-of-interest (VOI)-based analysis quantifying the reduction of specific bindings from baseline, which was homogenous across several brain regions within a blocking condition (S1 Fig). We defined receptor occupancy as the degree of reduction of specific binding using the values from striatal VOI, since they appeared to be the most reliable (see Methods) [40]. In 3 monkeys, we measured the relationship between D1R occupancy and the dose of SCH23390, which was approximated by a Hill function (Fig 1C; Eq. 4). We found that treatment with SCH23390 at doses of 100 and 30 μg/kg corresponded to 81% and 57% of D1R occupancy, respectively.

**Fig. 1.**
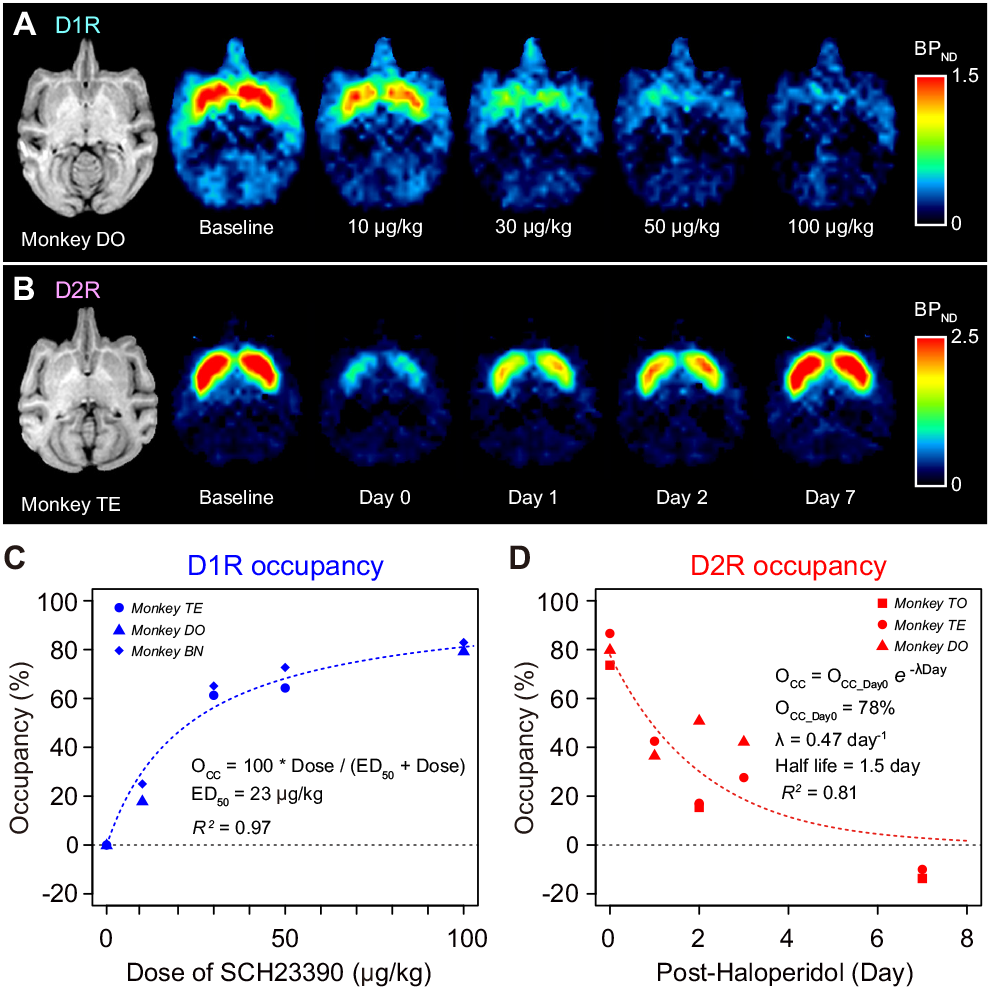
D1R and D2R occupancy measured by PET. (A) Representative horizontal MR (left) and parametric PET images showing specific b inding (BP_ND_) of [^11^C]SCH23390 at baseline and following drug-treatment with SCH23390 (10, 30, 50, or 100 μg/kg, i.m.). (B) Representative horizontal MR (left) and parametric PET images showing specific binding (BP_ND_) of [^11^C]raclopride at baseline and on 0 to 7 days after injection with haloperidol (10 μg/kg, i.m.). Color scale indicates BP_ND_ (regional binding potential relative to non-displaceable radioligand). (C) Occupancy of D1R measured at striatal ROI is plotted against the dose of SCH23390. (D) Occupancy of D2R measured at striatal ROI is plotted against the day after haloperidol injection. Dotted curves in C and D are the best fit o f E qs. 4 a nd 5, respectively. The data underlying this figure c an b e found o n t he following public repository: https://github.com/minamimoto-lab/2021-Hori-DAR.

Haloperidol was used for D2R antagonism. Unlike SCH23390, which was rapidly washed from the brain within a few hours, a single dose of haloperidol treatment was expected to show persistent D2R occupancy for the following several days as described in humans and mice [41, 42], providing the opportunity to test different occupancy conditions. The baseline [^11^C]raclopride PET image showed the highest radiotracer binding in the striatum (Fig 1B, Baseline). As expected, striatal binding was diminished not only just after pre-treatment with haloperidol (10 μg/kg, i.m.), but also on post-haloperidol day 2 (Fig 1B, Day 2). Binding had returned to the baseline level by day 7 (Fig 1B, Day 7). We measured D2R occupancy on days 0, 1, 2, 3, and 7 after a single haloperidol injection in 3 monkeys. An exponential decay function approximated the relationship between D2R occupancy and post-haloperidol days (Eq. 5); a single injection of haloperidol yielded 78% and 48% of D2R occupancy on days 0 and 1, respectively (Fig 1D).

### Effects of D1R- and D2R-blockade on behavior

We next quantified the effects of DAR blockade on behavior using a total of three monkeys not used in the PET occupancy study (monkeys KT, ST and MP; the former two for incentive and all three for cost-based motivation, respectively). Our goal here was to study the influence o f D 1R a nd D 2R manipulation on how monkeys adjusted their behavior based on expected benefits (reward size) or expected costs (delay or workload). We use two tasks where reward could be obtained by performing a simple action (releasing a bar). In each version of the task, we manipulated costs (delay or workload) or benefits (reward size), such that distinct trials corresponded to different levels of cost or benefits. A t t he b eginning of each trial, a visual cue provided information about the current cost and benefit, so that monkeys could adjust their behavior accordingly. We evaluated motivational processes by using computational modeling to measure the impact of incentive or costs on two behavioral measures: refusal rate (whether monkeys accepted or refused to perform the offered option; see below) and reaction time (RT; how quickly they respond).

### Effects of D1R- and D2R-blockade on benefit-based motivation

To assess the effect of blockade of D1R and D2R on benefit-based motivation, we tested 2 monkeys with a reward-size task (Fig 2A). In every trial of this task, the monkeys were required to release a bar when a visual target changed from red to green to get a liquid reward. A visual cue indicated the amount of reward (1, 2, 4, or 8 drops) at the beginning of each trial (Fig 2A). All monkeys had been trained to perform basic color discrimination trials in the cued multi-trial reward schedule task [43] for more than 3 months. As in previous experiments using a single option presentation, the action was very easy and monkeys could not fail if they actually tried to release the bar on time; the error rate is much lower in the absence of information about costs and benefits [2, 44]. As in those previous experiments manipulating information regarding costs and benefits, failures (either releasing the bar too early or too late) were usually observed in small reward trials and/or close to the end of daily sessions. Therefore, they were regarded as trials in which the monkeys refused to release the bar, presumably because they were not sufficiently motivated to correctly release the bar (i.e., refusal) [2]. Hence, the refusal rate provides a reliable measure of the influence of motivation on behavior [9, 45–48]. We previously showed that the refusal rate (*E*) is inversely related to reward size (*R*), which has been formulated with a single free parameter *a* [2] (Fig 2B),

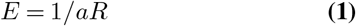

**Fig. 2.**
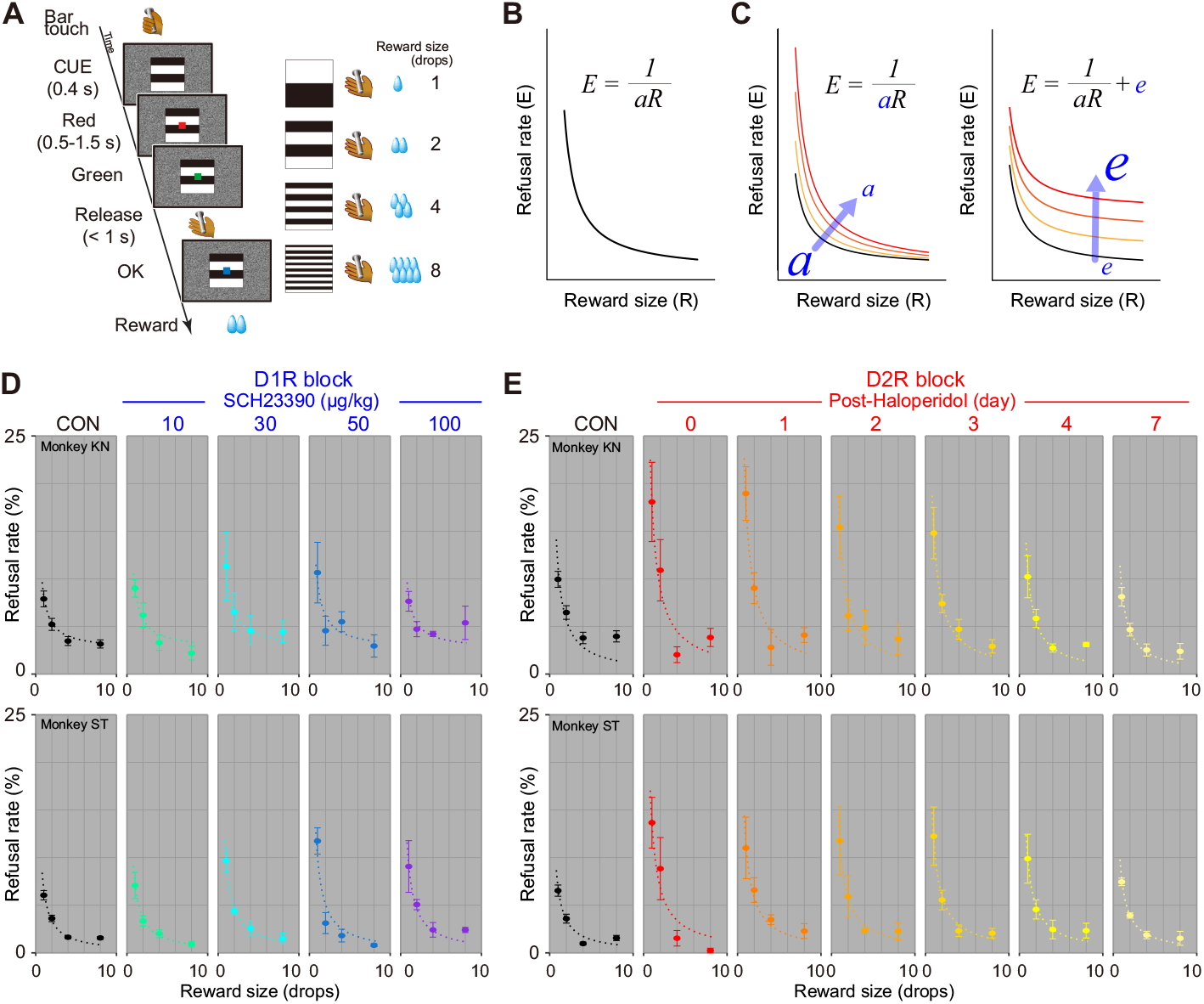
D1R/D2R blockade increased refusal rates in reward-size task. (A) Reward-size task. Left: Sequence of events during a trial. Right: Association between visual cues and reward size. (B) Schematic illustration of inverse function between refusal rate and reward size. (C) Schematic illustration of two explanatory models of decrease in motivation. Left: Increase in refusal rate (i.e., decrease in motivation) in relation to reward size caused by decrease in incentive impact (*a*). Right: An alternative model explaining increase in refusal rate irrespective of reward size. (D-E) Behavioral data under D1R and D2R blockade, respectively. CON, control. Refusal rates (mean ± SEM) as a function of reward size for monkeys KN (top) and ST (bottom). Dotted curves are the best-fit of inverse function (S1 Table). The data underlying this figure can be found on the following public repository: https://github.com/minamimoto-lab/2021-Hori-DAR.

In agreement with these previous studies, both monkeys exhibited the inverse relationship in non-treatment condition (Fig 2D and 2E, Control).

For D1R blockade, the monkeys were tested with the task 15 min after a systemic injection of SCH23390 (10, 30, 50, and 100 μg/kg) or vehicle as control. D1R blockade increased the refusal rates particularly in smaller reward size trials (Fig 2D). We considered whether this increase was due to a reduction in the incentive impact of reward, or a decrease in motivation irrespective of reward size. These factors can be captured by a decrease in parameter *a* of the inverse function and implementing intercept *e*, respectively (Fig 2C). To quantify the increases in refusal rate, we compared 4 models while considering these two factors as random effects: model #1, random effect on *a*; model #2, random effect on *a* with fixed *e*; model #3, fixed *a* with random effect on *e*; model #4, random effect on both *a* and *e* (see S1 Table). For both monkeys, the increases in refusal rate were explained by a decrease in the parameter *a* due to the treatment, while the inverse relation with reward size was maintained (model #3 for monkey KN and model #1 for ST; S1 Table). We then assessed changes in parameter *a*, which indicates the incentive impact of reward size. As shown Figure 3A, normalized *a* became smaller as the dose of SCH23390 was increased to 30 or 50 μg/kg, but then it increased at the highest dose (100 μg/kg) for monkeys KN but less clearly so for monkey ST (Fig 3A, left).

**Fig. 3.**
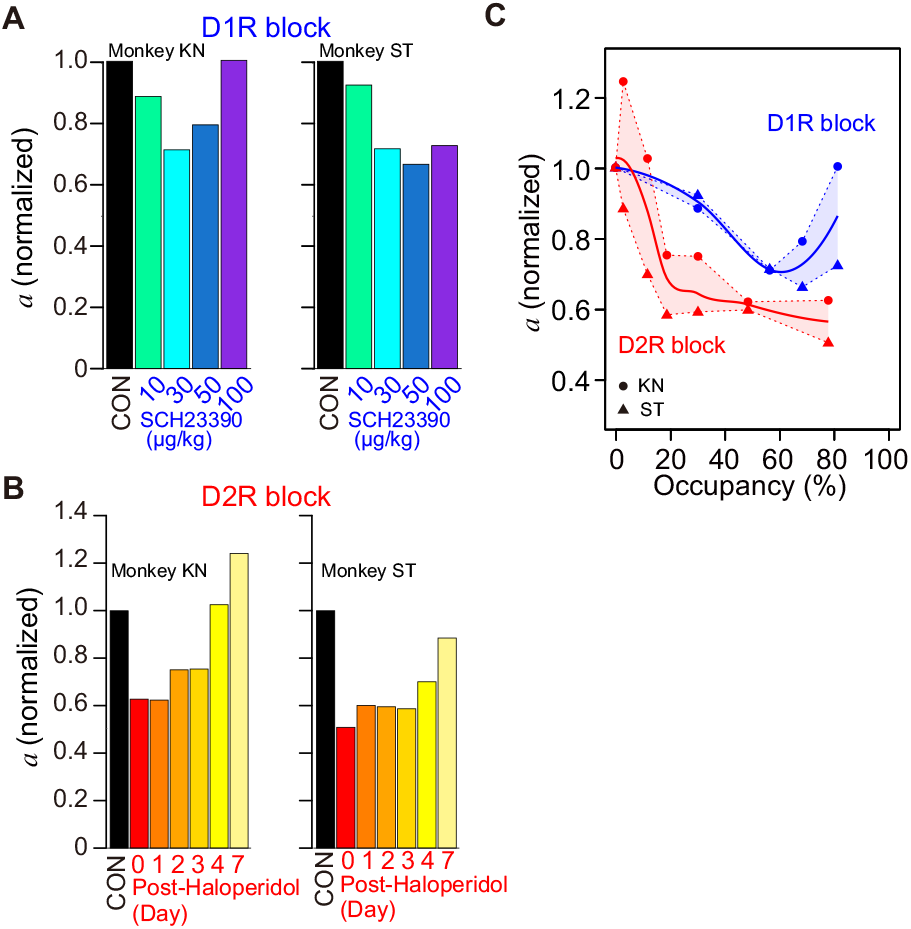
Effect of D1R/D2R blockade on incentive impact of reward size. (A) Bars indicate normalized incentive impact (*a*) for each treatment condition under D1R blockade for monkeys KN and ST. The value was normalized by the value of control condition. (B) Same as A, but for D2R blockade. (C) Relationship between an incentive impact and occupancy for D1R (blue) and D2R blockade (red). Thick curves indicate locally weighted smoothing (LOESS) of individual data (filled circles and triangles for monkeys KN and ST, respectively). The data underlying this figure can be found on the following public repository: https://github.com/minamimoto-lab/2021-Hori-DAR.

For D2R blockade, the monkeys were tested with the task 15 min after a single injection of haloperidol (10 μg/kg, i.m., day 0), and they were then successively tested on the following days 1, 2, 3, 4 and 7. We also found an increase in refusal rates for D2R blockade in both monkeys: the refusal rates were highest on the day of haloperidol injection, after which they decreased as the days went by (Fig 2E). Similar to the D1R blockade, the increases in refusal rate due to D2R blockade were explained solely by a decrease of parameter *a* according to the days following the treatment for both monkeys (Fig 2E, S1 Table; model #1 for both monkeys KN and ST). Our model-based analysis revealed that a decreased about 40% on the day of haloperidol injection and the following 3 days as compared to control, and then recovered to almost the control level by day 7 (Fig 3B).

To compare the effects between D1R and D2R blockades directly, we plotted changes in incentive impact along with the degree of blockage that was normalized across 3 monkeys (Fig 3C). In both D1R and D2R blockades, *a* declined according to the increase in occupancy; it gradually declined as D1R occupancy increased, but then increased at the highest occupancy, presenting a U-shaped tendency, whereas it steeply declined until 20% D2R occupancy, and then continued to decrease slightly until 80% occupancy (Fig 3C). At 20 – 80% occupancy, the incentive impacts for D2R blockade stayed lower than those for D1R, suggesting a stronger sensitivity of incentive impact to D2R blockade.

We sought to verify that the effect of D2R antagonism was not specific f or h aloperidol, a nd a lso t o v alidate t he comparison between D1R and D2R in terms of receptor occupancy. We examined the behavioral effect of another D2R antagonist, raclopride, at a dose yielding about 50% receptor occupancy (10 μg/kg, i.m.; S2 Fig). Following this dose of raclopride administration, a monkey again exhibited increased refusal rates, which was explained by inverse function with *a =* 5.2 (drop-1), a comparative value of incentive impact observed at 50% D2R occupancy with haloperidol [*a =* 5.4 (drop-1), day 1; S2B Fig]. Thus, our data suggest that D2R antagonism-induced reduction of the incentive effect seems to reflect the degree of receptor blockade regardless of the antagonist used.

### Effects of D1R- and D2R-blockade on response speed

To evaluate the extent to which the influence o f DAR manipulation in the reward size task could affect another behavioral measure through a single motivational process, we examined RT modulations across trials. Consistent with previous studies using systemic administration of D1R or D2R antagonists (e.g., [49]), DAR blockade in our study prolonged RTs in a treatment-dependent manner. For D1R blockade, RTs were increased according to the antagonist dose (2-way ANOVA, main effect of treatment, *p <* 1.0 × 10^−13^ for both monkeys; e.g., S3A-C Fig, see details in legend). D1R antagonism also tended to increase the proportion of late release (2-way ANOVA, main effect of treatment, *p =* 0.08, monkey KN; *p =* 0.0038 monkey ST; e.g., S3D Fig). A simple account of these effects of D1R manipulation on RT is that the modulations in RT across conditions are caused by changes in motivation, such that the positive impact of reward on behavior affects both whether monkeys perform the action (refusal rate) as well as how quickly they will respond (RT). We reasoned that, if this were the case, then the intersession variability in RT and refusal rate should be correlated. A session-by-session analysis revealed that there was indeed a significant l inear r elationship between refusal rates and RTs in both monkeys, even when the treatment conditions were changed (Fig 4A; S2 Table). D2R blockade also prolonged RTs (main effect of treatment, *p* < 1.0 × 10^−4^; e.g., S3E-G Fig). D2R blockade did not change the refusal patterns (i.e., too early or late release) (2-way ANOVA, treatment, *p =* 0.31; e.g., S3H Fig). As with the case of D1R, there was a linear relationship between refusal rates and RTs across D2R antagonism sessions, in which treatment had no discernible effect on the steepness of the slope (Fig 4B; S2 Table). Collectively, these results indicate that refusal rate and RT were similarly affected by DAR manipulation, in line with the concept that DAR affects a central process (motivation), which controls the influence of expected reward on both action selection and execution.

**Fig. 4.**
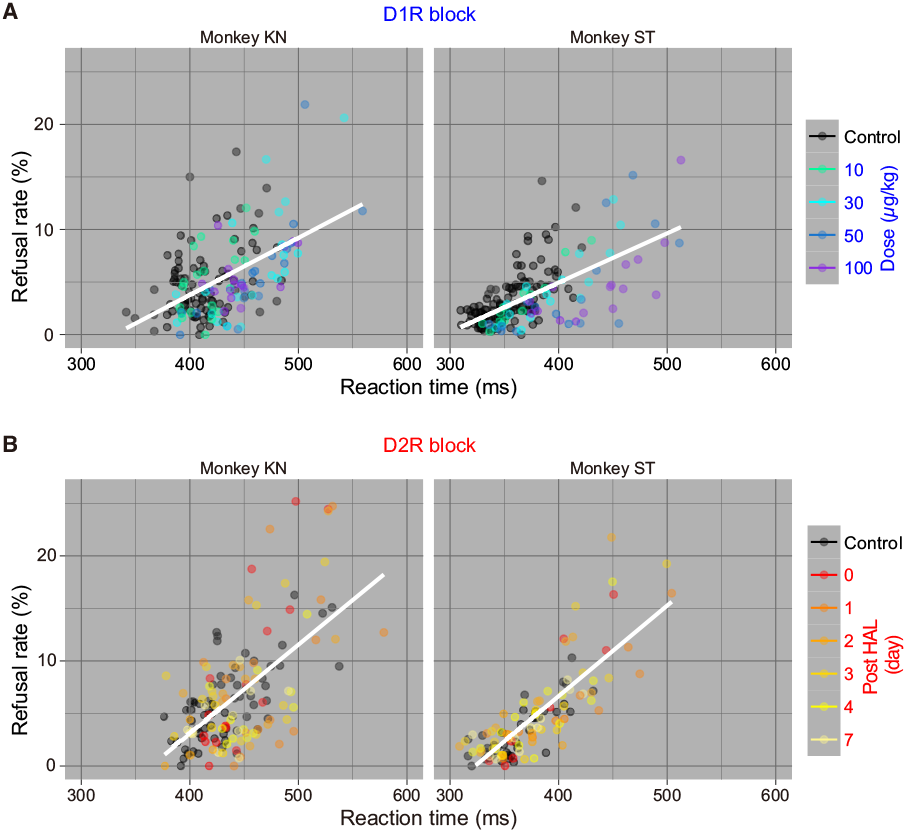
Relationship between refusal rate and reaction time in reward size task. (A) Relationship between refusal rate and average reaction time for each reward size in session-by-session for D1R blocking in monkeys KN and ST. Colors indicate treatment condition. (B) Same as A, but for D2R blocking. Note that a simple linear regression (white line, model #1 in S2 Table) was selected as the best-fit model to explain the d ata. The data underlying this figure can be found on the following public repository: https://github.com/minamimoto-lab/2021-Hori-DAR.

### Little influence of D1R- or D2R-blockade on hedonic impact of reward

The behavioral data shown above suggest that blockade of DAR attenuates the incentive effect of reward on behavior. To evaluate the impact of DAR blockade on other aspects of motivation and reward processing, we also examined the effect of DAR manipulation on hedonic processes, i.e., on how pleasant was the reward consumption. In line with previous experiments in rodents [15, 50], we did not find any effect of treatment with D1R or D2R antagonist on overall intake or sucrose preference in either of the 2 monkeys tested (S4A Fig; see legend). We also assessed blood osmolality, a physiological index of dehydration and thirst drive [51], before and after the preference test. Again, DAR treatment had no significant influence on overall osmolality or recovery of osmolality (rehydration) (S4B Fig; see legend). These results suggest that DAR blockade has no influence on hedonic impact of reward. These results also support the notion that the increased refusal rate was not directly due to a reduction of thirst drive. In short, these results indicate that both D1R and D2R are involved in incentive motivation, i.e., in the positive influence of the expected reward size on behavior (refusal rate and RT), but not in the hedonic impact of reward. We next examined the influence of DAR manipulation on cost processing.

### Differential effects of D1R and D2R blockades on workload and delay discounting

The trade-off between the reward and costs of obtaining the reward affects decision-making as well as motivation. Both humans and animals have the tendency to prefer immediate, smaller rewards over larger, but delayed rewards. The preference can be predicted by discounting the reward’s intrinsic value by the duration of the expected delay, an effect designated as “delay discounting” [52, 53]. Discounting of the reward value also occurs in proportion to the predicted effort needed to obtain the rewards; an effect called “effort discounting” [54]. Delay and effort discounting are typically measured in choice tasks, providing the relative impact of costs on reward in decision-making. Previously, we measured the discounting effect of these costs on outcome value by quantifying the relation between the amount of expected cost and the change in operant, reward-directed behavior [55].

In this study, we used the same procedure to assess the effect of selective DAR blockade on cost-based motivation. For this purpose, we used a work/delay task (Fig 5A), where the basic features were the same as those in reward-size task. There were two trial types. In the work trials, the monkeys had to perform 0, 1, or 2 additional instrumental actions to obtain a fixed amount of reward, and the cost (workload) scaled with the number of trials to perform. In the delay trials, after the monkeys correctly performed one instrumental trial, a reward was delivered 0–7 seconds later, such that the cost (delay) scaled with the time between action and reward delivery. Note that here, as in most natural conditions, greater workload is inherently associated with longer delays. Thus, in an attempt to isolate the effort component, we adjusted the delay for reward in delay trials based on the duration of corresponding workload trials: since the timing of the trials is matched between workload and delay trials, they only differed in the number of actions and therefore in the amount of effort. At the beginning of each trial, the cost (workload or delay) was indicated by a visual cue that lasted throughout the trial. As with the reward task, we used computational modeling to quantify the influence of cost information on behavior. We have shown that the monkeys exhibited linear relationships between refusal rate (*E*) and remaining costs (*CU*) for both work and delay trials, as follows:

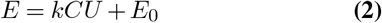

where *k* is a coefficient and *E_0_* is an intercept [43] (Fig 4B). By extending the inference and formulation of reward-size task (Eq. 1), this linear effect proposes that the reward value is hyperbolically discounted by cost, where the coefficient *k* corresponds to discounting factors.

**Fig. 5.**
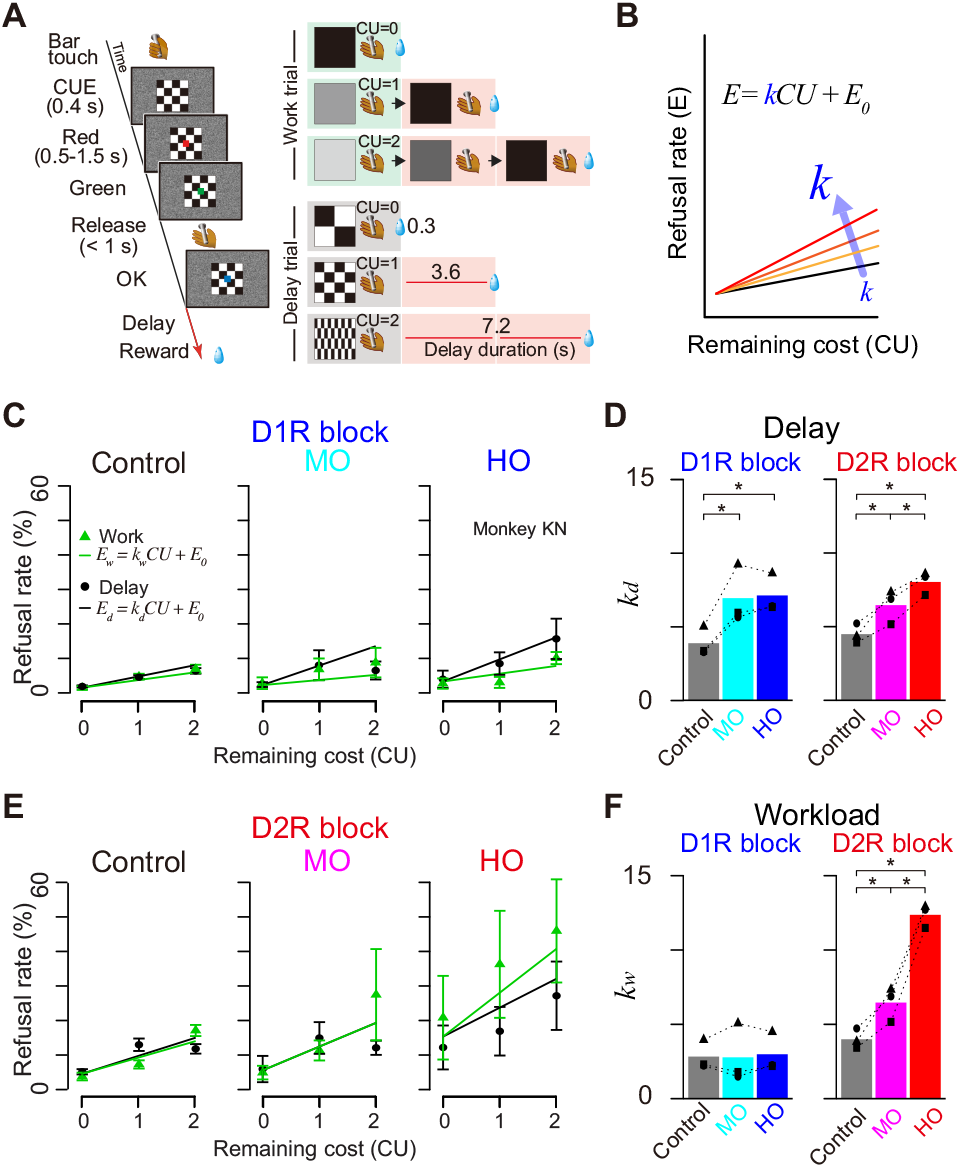
Differential effects of D1R and D2R blockade on cost-based motivational valuation. (A) The work/delay task. The sequence of events (left) and relationships between visual cues and trial schedule in the work trials (right top 3 rows) or delay duration in the delay trials (right bottom 3 rows) are shown. *CU* denotes the remaining (arbitrary) cost unit to get a reward, i.e., either remaining workload to perform trial(s) or remaining delay periods. (B) Schematic illustration of an explanatory model of increases in refusal rate by increasing cost sensitivity (*k*). (C) Effects of D1R blockade. Representative relationships between refusal rates (monkey KN; mean ± SEM) and remaining costs for workload (green) and delay trials (black). Saline control (Control), moderate (30 μg/kg; MO) and high D1R occupancy treatment condition (100 μg/kg; HO) are shown. Green and black lines are the best-fit lines for work and delay trials in model #1 in S3 Table, respectively. (D) Effects of D2R blockade. Non-treatment control (Control), moderate (1 day after haloperidol; MO) and high D2 occupancy treatment conditions (day of haloperidol; HO) are shown. Others are the same for C. (E) Comparison of effects between D1R and D2R blockade on delay-discounting parameter (*kd*). Bars and symbols indicate mean and individual data, respectively. (F) Comparison of effects between D1R and D2R blockade on workload-discounting parameter (*kw*). Asterisks represent significant difference (* *p <* 0.05, one-way ANOVA with post-hoc Tukey HSD test). The data underlying this figure can be found on the following public repository: https://github.com/minamimoto-lab/2021-Hori-DAR.

We tested 3 monkeys (monkeys KN, MP, and ST) and obtained a refusal rate to infer delay- and workload-discounting. We confirmed that refusal rates of control condition increased as the remaining cost increased (e.g., Fig 5C, control). Figure 5B illustrates our hypothesis that DAR blockade increases cost sensitivity (i.e., discounting factor, *k*), thereby elevating the proportion of an increase in refusal rate relative to remaining cost.

To compare the effect of D1R vs D2R antagonism on cost sensitivity at the same degree of receptor blockade, we assessed the performance of the monkeys under two comparable levels of DAR occupancy for D1R and D2R at about 50% (called “moderate occupancy” or MO) and 80% (“high occupancy” or HO), and under baseline condition (non-treatment) as control. According to the occupancy study (Fig 1), MO and HO conditions corresponded to pre-treatment with 30 and 100 μg/kg of SCH23390 for D1R, and 1 day after and the day of haloperidol treatment for D2R, respectively. Linear mixed models (LMM) analysis verified the assumption that DAR blockade increased delay and workload discounting independently without considering the random effect of treatment condition or subject (Fig. 5CD, S3 Table; see Methods). We found that delay discounting was significantly increased according to the degree of DAR blockade irrespective of receptor subtype (D1, *F*(2, 4) = 36.9, *p =* 0.0026; D2, *F*(2, 4) = 41.4, *p =* 0.0021; Fig 5E). Workload-discounting (*kw*), on the other hand, was specifically increased by D2R blockade in an occupancy-dependent manner (one-way ANOVA, main effect of occupancy; D1, *F*(2, 4) = 0.125, *p =* 0.89; D2, *F*(2, 4) = 243.2, *p =* 6.6 × 10-5; Fig 5F).

In line with what we found in the reward size task, D1R blockade did not have any significant effect on the linear relation between refusal rate and reaction time in either trial type (Fig 6A). Thus, the influence of D1R manipulation on behavior could readily be accounted for by a single variable, which affects both RT and refusal rate. By contrast, D2R blockade produced an occupancy dependent increase in the steepness of the linear relation between RT and refusal rate in workload trials, but not in delay trials (Fig 6B). Thus, D2R manipulation had a distinct influence on RT and refusal rate in workload trials, suggesting that it was acting on behavior through a distinct motivational process such as overcoming effort costs (see Discussion).

**Fig. 6.**
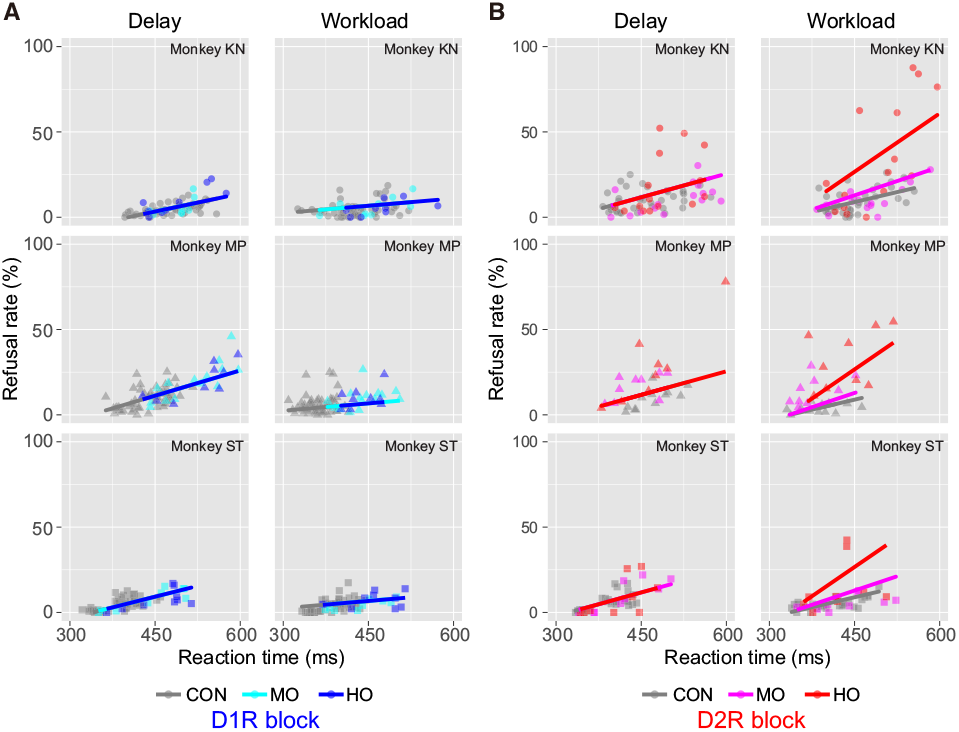
Relationship between refusal rate and reaction time in work/delay task. (A) Relationship between refusal rate and average reaction time for each remaining cost in session-by-session for D1R blocking in delay and workload trials. Data are plotted individually for monkeys KN, MP, and ST, in order from top to bottom. Colors indicate treatment condition. Thick lines indicate linear regression lines. (B) Same as A, but for D2R blocking. Note that for the data in workload trials under D2R treatment, a linear model with random effect of condition (model #4 in S4 Table) was chosen as the best model to explain the data, whereas for the other data, a simple linear regression model (model #1, without any random effect or model #2 with random effect of subject) was selected. The data underlying this figure can be found on the following public repository: https://github.com/minamimoto-lab/2021-Hori-DAR.

### Joint influences of D1R and D2R blockades on motivation

Considering the direct and indirect striatal output pathways where neurons exclusively express D1R and D2R, respectively, and the potential functional opposition between these pathways [56], we examined the effect of joint blockade of D1R and D2R. To facilitate the comparison of the influence of two receptors, we examined the behavioral effects of both D1R and D2R blockades at the same occupancy level. After treatment with both SCH23390 (100 μg/kg) and haloperidol (10 μg/kg), seemingly achieving ~80% of occupancy for both subtypes (cf. Fig 1C and 1D), all monkeys stopped performing the task with a small number of correct trials (1-13% of control). When we treated the monkeys with SCH23390 (30 μg/kg) on the day following that of haloperidol injection (i.e., both D1R and D2R assumed to be occupied at ~50%), the monkeys had higher refusal rates in delay trials than control (Fig 7A, D1R+D2R block), such that discounting factor (*kd*) became significantly h igher t han in control conditions (*p <* 0.05, Tukey HSD test; Fig 7B, delay). By contrast, this simultaneous D1R and D2R blockade appeared to attenuate the effect of D2R antagonism on workload: the refusal rates in work trials were not as high as in D2R blockade alone (Fig 7A), and the difference in workload-discounting factor (*kw*) between treated and control or baseline conditions disappeared (*p >* 0.05; Fig 7B, workload). A similar tendency of counterbalancing influence was also seen in the motivation for minimum cost trials (*E_0_*) (Fig 7B). These results suggest that blocking both receptor subtypes tends to induce a synergistic effect on delay-discounting, while their effects on workload-discounting cancel each other out.

**Fig. 7.**
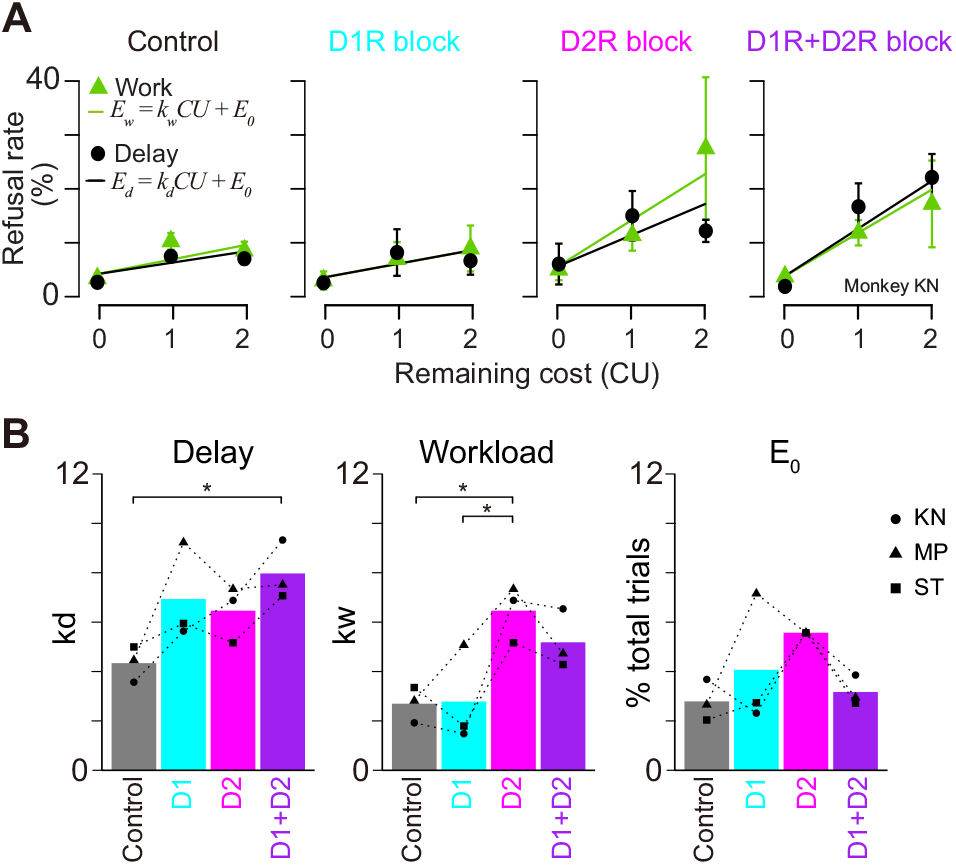
Effect of both D1R and D2R blockades on cost evaluation for motivation. (A) Representative relationship between refusal rates (in monkey KN; mean ± SEM) and remaining costs for workload (green) and delay trials (black). Best-fit parameters, workload-discounting (*kw*), delay-discounting (*kd*), and intercept (E0), are plotted for each treatment condition. Bars and symbols indicate mean and individual data, respectively. D1R+D2R indicates the data obtained under both D1R and D2R blockades at moderate occupancy, while D1R and D2R blockades at high occupancy resulted in almost no correct performance (see text). All parameters are derived from the best fit of model #1 in S3 Table. Asterisks represent significant difference (\**p <* 0 .05, one-way ANOVA with post-hoc Tukey HSD test). The data underlying this figure can be found on the following public repository: https://github.com/minamimoto-lab/2021-Hori-DAR.

Finally, since workload trials revealed a potential specific action of D2R, with a dissociation between refusal rate and RT effects, we examined the joint influence of D1R and D2R manipulations. As shown in S5B Fig., when D1R and D2R were simultaneously blocked, the relationship between RT and refusal rate in work trials became closer to that of control monkeys than those treated with D2R agonists alone, consistent with their impacts on refusal rate (cf. Fig 7B). Therefore, even if D1R antagonist alone had little effect on workload sensitivity, it may be able to counteract the effect of D2R treatment under these conditions.

## Discussion

Combining the PET occupancy study and pharmacological manipulation of D1- and D2-like receptors with quantitative measurement of motivation in monkeys, the current study demonstrated dissociable roles of the DA transmissions via D1R and D2R in the computation of the cost/benefits trade-off to guide action. To the best of our knowledge, this is the first study to directly compare the contribution of dopamine D1R and D2R along with the degree of receptor blockade. Using model-based analysis, we showed that DAR blockade had a clear quantitative effect on the sensitivity of animals to information about potential costs and benefits, without any qualitative effect on the way monkeys integrated costs and benefits and adjusted their behavior. We showed that blockade of D1R or D2R reduced the incentive impact of reward as the degree of DAR blockade increased, and the incentive impact was more sensitive to the D2R blockade than the D1R blockade at lower occupancy. In cost-discounting experiments, we could dissociate the relation between each DAR type and workload vs delay-discounting: workload-discounting was increased exclusively by D2R antagonism, whereas delay-discounting was increased by DAR blockade irrespective of receptor subtype. When both D1R and D2R were blocked simultaneously, the effects were synergistic and strengthened for delay-discounting, while the effects were antagonistic and diminished for workload-discounting. These results suggest that the mechanism in the action of DA is different for incentive motivation and temporal discounting, and for workload discounting.

### DA controls the incentive effect of expected reward amount

Previous pharmacological studies have shown that DAR blockade decreased the speed of action and/or probability of engagement behavior [22, 23]. However, the previous studies did not measure the effect of DAR blockade on incentive motivation in multiple rewarding conditions, and therefore data describing the quantitative relationship among DAR stimulation, reward, and motivation are not available. In the present study, we used a behavioral paradigm that enabled us to formulate and quantify the relationship between reward and motivation [2] (Fig 2). Our finding, a reduction of incentive impact due to DAR antagonism (cf., Fig 3) is in line with the incentive salience theory, that is, DA transmission attributes salience to incentive cue to promote goal-directed action [12]. The lack of effect of DA manipulation on satiety and spontaneous water consumption is consistent with previous studies in rodents [57, 58]. Our results are also compatible with the idea that DA manipulation mainly influences incentive processes (influence of reward on action) but does not cause a general change of reward processing, which includes hedonic processes (evaluation itself, pleasure associated with consuming reward) [59, 60], although further experiments would be necessary to address that point directly [12].

Our model-based analysis indicates that DAR blockade only had a quantitative influence (reduction of incentive impact of reward) without changing the qualitative relationship between reward size and behavior. This is in marked contrast to the reported effects of inactivation of brain areas receiving massive DA inputs, including the orbitofrontal cortex, rostromedial caudate nucleus, and ventral pallidum. Indeed, in experiments using nearly identical tasks and analysis, inactivation or ablation of these regions produced a qualitative change in the relationship between reward size and behavior (more specifically, a violation of the inverse relationship between reward size and refusal rates) [47, 48, 61]. Thus, the influence of DAR cannot be understood as a simple permissive or activating effect on target regions. The specificity of the DAR functional role is further supported by the subtle, but significant difference between the behavioral consequences of blocking of D1R vs D2R. By combining a direct measure of DAR occupancy and quantitative behavioral assessment, the present study demonstrates that the incentive impact of reward is more sensitive to D2R blockade than D1R blockade, and especially at a lower degree of occupancy (cf. Fig 3C). Moreover, the relationship between occupancy and incentive impact was monotonous for D2R, but tended to be U-shaped for D1R. Although this U-shaped effect of D1R blockade was inferred solely based on the refusal rate of two monkeys without statistical support at the population level and was not found in the RT data, such non-monotonic effects have been repeatedly reported. For example, working memory performance and related neural activity in the pre-frontal cortex takes the form of an “inverted-U” shaped curve, where too little or too much D1R activation impairs cognitive performance [62–64]. As for the mechanisms underlying the distinct functional relation between the behavioral effects of D1R vs D2R blockade, it is tempting to speculate that this is related to a difference in their distribution, their affinity and the resulting relation with phasic vs tonic DA action. Indeed, DA affinity for D2R is 100 times higher than that for D1R [65]. This is directly in line with the stronger effect of D2R antagonists at low occupancy levels. Moreover, in the striatum, a basal DA concentration of 5–10 nM is sufficient to constantly stimulate D2R. Using available biological data, a recent simulation study showed that the striatal DA concentration produced by the tonic activity of DA neurons (40 nM) would occupy 75% of D2R but only 3.5% of D1R [66]. Thus, blockade of D2R at low occupancy may interfere with tonic DA signaling, whereas D1R occupancy would only be related to phasic DA action, i.e., when transient but massive DA release occurs (e.g., in response to critical information about reward). We acknowledge that this remains very hypothetical, but irrespective of the underlying mechanisms, our data clearly support the idea that DA action on D1R vs D2R exerts distinct actions on their multiple targets to enhance incentive motivation.

### DA transmission via D1R and D2R distinctively controls cost-based motivational process

Although many rodent studies have demonstrated that attenuation of DA transmission alters not only benefit but also cost-related decision-making, the exact contribution of D1R and D2R remains elusive. For example, reduced willingness to exert physical effort to receive higher reward was similarly found following D1R and D2R antagonism in many rodent studies [24, 26, 59], while it was observed exclusively by D2 antagonism in other studies [21, 25]. This inconsistency may have two reasons. First, previous studies usually investigated the effect of antagonism on D1R and D2R along with a relative pharmacological concentration (e.g., low and high doses). In the present study, PET-assessed DAR manipulation allowed us to directly compare the behavioral effect between D1R and D2R with an objective reference, namely occupancy (i.e., 50% and 80% occupancy). Second, the exact nature of the cost (effort vs delay) has sometimes been difficult t o identify, and effort manipulation is often strongly correlated with reward manipulation (typically when the amount of reward earned is instrumentally related to the amount of effort exerted, see [10]). Here, using a task manipulating forthcoming workload independently from reward value, we demonstrated that blockade of D2R, but not D1R, increased workload-discounting in an occupancy-dependent manner while maintaining linearity (cf. Fig 5F).

Delay-discounting and impulsivity — the tendency associated with excessive delay-discounting — are also thought to be linked to the DA system [69, 70]. Systemic administration of D1R or D2R antagonist increases preference for immediate small rewards, rather than larger and delayed rewards [25, 32–34]. Concurrently, some of these studies also showed negative effects of D1R [34] or D2R blockade [33] on impulsivity. These inconsistencies may be attributed to the differences in behavioral paradigms or drugs (and doses) used. Our PET-assessed DAR manipulation demonstrated that blockade of D1R and D2R at the same occupancy level (50% and 80%) similarly increased delay-discounting (Fig 5E), suggesting that DA transmission continuously adjusts delay-discounting at the post-synaptic site. This observation is in good accord with the previous finding that increasing DA transmission decreases temporal discounting; e.g., amphetamine or methylphenidate increased the tendency to choose long-delays options for larger rewards [32–34, 71, 72].

In sharp contrast to incentive motivation and delay discounting, which involve both D1R and D2R, the following three observations illustrate a unique mechanism of DA action on workload discounting through D2R only. First, workload-discounting increased only with D2R antagonism (cf. Fig 5F). Second, the occupancy-dependent effect of D2R antagonism on the decision—response time relationship was only seen in workload trials (cf. Fig 6). Third, D1R and D2R had a synergistic effect in the delay-discounting trials, but an antagonistic effect in the workload-discounting trials (cf. Fig 7). These results extend previous studies demonstrating increased effort-discounting by D2R blockade [25, 73]. Besides, our observation that blocking D2R increased refusal rates without slowing response (cf. Fig 6B) empathizes the role of DA in effort-based decision-making, and supports the notion that DA activation allows overcoming effort costs [21]. This is in apparent contrast to neurophysiological and voltammetry studies that show a lack of sensitivity of DA release to effort, but comforts the idea that DA function requires the integration of receptor action on top of neuronal activity and releasing patterns [10, 17].

This differential relation between DA and delay vs workload might be related to the differential expression of these receptors in the direct vs indirect striatopallidal pathway, where the striatal neurons exclusively express D1R and D2R, respectively [74]. Opposing functions between these pathways have been proposed: activity of the direct pathway (D1R) neurons reflects positive rewarding events promoting movement, whereas activity of the indirect pathway (D2R) neurons is related to negative values mediating aversion or inhibiting movements [56, 75, 76] (but see [77]). DA increases the excitability of direct-pathway neurons, and this effect was reduced by D1R antagonism, decreasing motor output. DA reduces the responsiveness of indirect pathway neurons via D2R [74], and blockade of D2R would increase the activity, reducing motor output via decreased thalamocortical drive [78]. Interestingly, a neural network model has been proposed by considering these opposing DA functions of direct/indirect circuit embedded in reinforcement learning framework, successfully explaining the enhancement of effort cost due to D2R blockade [79]. This scenario might also explain our finding of a synergistic effect of simultaneous D1R and D2R blockade on delay-discounting. Further work would be necessary to clarify this hypothesis, including the dynamic relation with tonic vs phasic DA release, but altogether, these data strongly support the concept that distinct neurobiological processes underlie benefits (reward availability) and costs (energy expenditure).

### Limitations of the current study

Finally, limitations of the current study and areas for further research can be discussed. First, there were relatively larger individual disparities in estimated values of D2R occupancy by haloperidol in three monkeys (cf. Fig 1D), which could reflect individual variance of haloperidol metabolism and/or elimination. However, the time course of the recovery of D2R occupancy was relatively consistent across subjects, being in line with that of behavioral change. Haloperidol induced long-lasting occupancy for several days, thereby potentially causing unexpected long-term changes, such as synaptic plasticity. Although we cannot eliminate the potential effects of plastic change, a comparable behavioral impact was also observed after raclopride administration, which would induce short-term occupancy (cf. S2 Fig), supporting the view that blockade of DAR reduced motivation in an occupancy dependent manner. Second, because of applying systemic antagonist administration, the current study could not determine which brain area(s) is responsible for antagonist-induced alterations of benefit- and cost-based motivation. While our data support the notion that differential neural networks involve workload-and delay-discounting, further study (e.g., local infusion of DA antagonist) is needed to identify the locus of the effects, generalizing our findings to unravel the circuit and molecular mechanism of motivation. We should also note that the current study does not address dynamic learning paradigms and therefore does not generalize our findings to the function of the DA system in learning directly. Despite these limitations, the current study provides unique insights into the role of the DA system in the motivational process.

## Conclusions

In summary, the present study demonstrates a dissociation between the functional role of DA transmission via D1- and D2-like receptors in benefit- and cost-based motivational processing. DA transmissions via D1R and D2R modulate both the incentive impact of reward size and the negative influence of delay. By contrast, workload-discounting is regulated exclusively via D2R, since apparently D1R alone had no role. In addition, D1R and D2R had synergistic roles in delay-discounting but opposite roles in workload-discounting. These dissociations indicate different underlying mechanisms of DA on motivation, which can be attributed to differential involvement of the direct and indirect striatofugal pathways. Together, our findings add an important aspect to our current knowledge concerning the role of DA signaling motivation based on the trade-off between costs and benefits, thus providing an advanced framework for understanding the pathophysiology of psychiatric disorders.

## Materials and Methods

### Ethics statement

All surgical and experimental procedures were approved by the Animal Care and Use Committee of the National Institutes for Quantum and Radiological Science and Technology (#09-1035), and were in accordance with the Institute of Laboratory Animal Research Guide for the Care and Use of Laboratory Animals.

### Subjects

A total of nine male adult macaque monkeys (8 Rhesus and 1 Japanese; 4.6-7.7 kg) were used in this study. All monkeys were individually housed. Food was available ad libitum, and motivation was controlled by restricting access to fluid to experimental sessions, when water was delivered as a reward for performing the task. Animals received water supplementation whenever necessary (e.g., if they could not obtain enough water during experiments), and they had free access to water whenever testing was interrupted for more than a week. For environmental enrichment, play objects and/or small foods (fruits, nuts, and vegetables) were provided daily in the home cage.

### Drug treatment

All experiments in this study were carried out with injected intramuscular (i.m.) SCH23390 (Sigma-Aldrich), haloperidol (Dainippon Sumitomo Pharma, Japan), and raclopride (Sigma-Aldrich) dissolved or diluted in 0.9% saline solution. Animals were pretreated with an injection of SCH23390 (10, 30, 50, or 100 μg/kg), haloperidol (10 μg/kg), or raclopride (10 or 30 μg/kg) 15 min before the beginning of behavioral testing or PET scan. In behavioral testing, saline was injected as a vehicle control by the same procedure as the drug treatment. The administered volume was 1 mL across all experiments with each monkey.

### Surgery

Four monkeys underwent surgery to implant a head-hold device for the PET study using aseptic techniques [80]. We monitored body temperature, heart rate, SpO2 and tidal CO2 throughout all surgical procedures. Monkeys were immobilized by i.m. injection of ketamine (5–10 mg per kg) and xylazine (0.2–0.5 mg per kg) and intubated with an endotracheal tube. Anesthesia was maintained with isoflurane (1–3%, to effect). The head-hold device was secured with plastic screws and dental cement over the skull. After surgery, prophylactic antibiotics and analgesics were administered. The monkeys were habituated to sit in a primate chair with their heads fixed for approximately 30 min for more than 2 weeks.

### PET procedure and occupancy measurement

Four monkeys were used in the measurement. PET measurements were performed with two PET ligands: [^11^C]SCH23390 (for studying D1R binding) and [^11^C]raclopride (for studying D2R binding). The injected radioactivities of [^11^C]SCH23390 and [^11^C]raclopride were 91.7 ± 6.0 MBq (mean ± SD) and 87.0 ± 4.9 MBq, respectively. Specific radioactivities of [^11^C]SCH23390 and [^11^C]raclopride at the time of injection were 86.2 ± 40.6 GBq/μmol and 138.2 ± 70.1 GBq/μmol, respectively. All PET scans were performed using an SHR-7700 PET scanner (Hamamatsu Photonics Inc., Japan) under conscious conditions and seated in a chair. Prior to the PET study, the monkeys underwent surgery to implant a head-hold device using aseptic techniques [65]. After transmission scans for attenuation correction using a 68Ge–68Ga source, a dynamic scan in three-dimensional (3D) acquisition mode was performed for 60 min ([^11^C]SCH23390) or 90 min ([^11^C]raclopride). The ligands were injected via crural vein as a single bolus at the start of the scan. All emission data were reconstructed with a 4.0-mm Colsher filter. Tissue r adioactive concentrations were obtained from volumes of interest (VOIs) placed on several brain regions where DARs are relatively abundant: caudate nucleus, putamen, nucleus accumbens (NAcc), thalamus, hippocampus, amygdala, parietal cortex, principal sulcus (PS), dorsolateral prefrontal cortex (dlPFC), and ventrolateral prefrontal cortex (vlPFC), as well as the cerebellum (as reference region). Each VOI was defined on individual T1-weighted axial magnetic resonance (MR) images (EXCELART/VG Pianissimo at 1.0 tesla, Toshiba, Japan) that were co-registered with PET images using PMOD® image analysis software (PMOD Technologies Ltd, Switzerland). Regional radioactivity of each VOI was calculated for each frame and plotted against time. Regional binding potentials relative to non-displaceable radioligands (BP_ND_) of D1R and D2R were estimated with a simplified reference tissue model on VOI and voxel-by-voxel bases [81–83]. The monkeys were scanned with and without drug-treatment condition on different days.

Occupancy levels were determined from the degree of reduction (%) of BP_ND_ by antagonists [69]. DA receptor occupancy was estimated as follows:

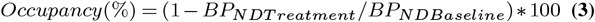

where BP_ND Baseline_ and BP_ND Treatment_ are BP _ND_ measured without (baseline) and with an antagonist, respectively. Relationship between D1R occupancy (D1 Occ) and dose of SCH23390 (Dose) was estimated with 50% effective dose (ED_50_) as follows:

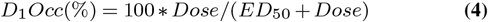

Relationship between D2R occupancy (D2Occ) and days after haloperidol injection was estimated using the level at day 0 with a decay constant (λ) as follows:

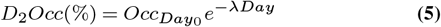

### Behavioral tasks and testing procedures

Three monkeys (ST, 6.4 kg; KN, 6.3 kg; M7, 7.3 kg) were used for the behavioral study. For all behavioral training and testing, each monkey sat in a primate chair inside a sound-attenuated dark room. Visual stimuli were presented on a computer video monitor in front of the monkey. Behavioral control and data acquisition were performed using the REX program. Neurobehavioral Systems Presentation software was used to display visual stimuli (Neurobehavioral Systems). We used two types of behavioral tasks, reward-size task and work/delay task, as described previously [2, 43]. Both tasks consisted of color discrimination trials (see Figs 2A and 5A). Each trial began when the monkey touched a bar mounted at the front of the chair. The monkey was required to release the bar between 200 and 1,000 ms after a red spot (wait signal) turned green (go signal). On correctly performed trials, the spot then turned blue (correct signal). A visual cue was presented at the beginning of each color discrimination trial (500 ms before the red spot appearing). In the reward-size task, a reward of 1, 2, 4, or 8 drops of water (1 drop = ~0.1 mL) was delivered immediately after the blue signal. Each reward size was selected randomly with equal probability. The visual cue presented at the beginning of the trial indicated the number of drops for the reward (Fig 2A). In the work/delay task, a water reward (~0.25 mL) was delivered after each correct signal immediately or after an additional 1 or 2 instrumental trials (work trial), or after a delay period (delay trials). The visual cue indicated the combination of the trial type and requirement to obtain a reward (Fig 4A). Pattern cues indicated the delay trials with the timing of reward delivery after a correct performance: either immediately (0.3 s, 0.2 – 0.4 s; mean, range), a short delay (3.6 s, 3.0 – 4.2 s), or a long delay (7.2 s, 6.0 – 8.4 s). Grayscale cues indicated work trials with the number of trials the monkey would have to perform to obtain a reward. We set the delay durations to be equivalent to the duration for 1 or 2 trials of color discrimination trials, so that we could directly compare the cost of 1 or 2 arbitrary units (cost unit; *CU*).

If the monkey released the bar before the green target appeared or within 200 ms after the green target appeared or failed to respond within 1 s after the green target appeared, we regarded the trial as a “refusal trial”; all visual stimuli disappeared, the trial was terminated immediately, and after the 1-s inter-trial interval, the trial was repeated. Our behavioral measurement for the motivational value of outcome was the proportion of refusal trials. Before each testing session, the monkeys were subject to ~22 hours of water restriction in their home cage. Each session continued until the monkey would no longer initiate a new trial (usually less than 100 min).

Before this experiment, all monkeys had been trained to perform color discrimination trials in the cued multi-trial reward schedule task for more than 3 months. The monkeys were tested with the work/delay task for 1-2 daily sessions as training to become familiar with the cueing condition. Each monkey was tested from Monday to Friday. Treatment with SCH23390 was performed every four or five d ays. On other days without SCH23390, sessions with saline (1 mL) treatment were analyzed as control sessions. Haloperidol was given every two or three weeks on Monday or Tuesday, because D2R occupancy persisted for several days after a single dose of haloperidol treatment (Fig 1D). The days before haloperidol treatment were analyzed as control sessions. Each dose of SCH23390 or a single dose of haloperidol was tested 4 or 5 times with the reward-size task and at least 3 times with the work/delay task per each animal.

### Sucrose preference test

Two monkeys (RO, 5.8kg; KY, 5.6kg) were used for the sucrose preference test. The test was performed in their home cages once a week. In advance of the test, water access was prevented for 22 h. The monkeys were injected with SCH23390 (30 μg/kg), haloperidol (10 μg/kg), or saline 15 min before the sucrose preference test. Two bottles containing either 1.5% sucrose solution or tap water were set into bottle holders in the home cage and the monkeys were allowed to freely consume fluids for 2h. The total amount of sucrose (*SW*) and tap water (*TW*) intake was measured and calculated as sucrose preference index (*SP*) as follows: *SP = (SW – TW) / (SW + TW)*. The position of sucrose and tap water bottles (right or left toward the front panel of the home cage) was counterbalanced across sessions and monkeys. Drugs or saline was injected alternatively once a week. We also measured the osmolality level in blood samples (1 mL) obtained immediately before and after each testing session.

### Behavioral data analysis

All data and statistical analyses were performed using the R statistical computing environment (R Development Core Team, 2004). The average error rate for each trial type was calculated for each daily session, with the error rates in each trial type being defined as the number of error trials divided by the total number of trials of that given type. The monkeys sometimes made many errors at the beginning of the daily session, probably due to high motivation/impatience; we excluded the data until the 1st successful trial in these cases. A trial was considered an error trial if the monkey released the bar either before or within 200 ms after the appearance of the green target (early release) or failed to respond within 1 s after the green target (late release). We did not distinguish between the two types of errors and used their sum except for the error pattern analysis. We performed repeated-measures ANOVAs to test the effect of treatment × reward size (for data in reward-size task) on reaction time, on late release rate (i.e., error pattern). Post-hoc comparisons were performed using Tukey HSD test, and a priori statistical significance was set at = 0.05.

We used the refusal rates to estimate the level of motivation because the refusal rates of these tasks (*E*) are inversely related to the value for action [2]. In the reward-size task, we used the inverse function (Eq. 1). We fitted the data to linear mixed models (LMMs) [85], in which the random effects across DAR blockade conditions on parameter *a* and/or intercept *e* (Fig 2C) were nested. Model selection was based on Bayesian information criterion (BIC), an estimator of in-sample prediction error for the nested models (S1 Table). Using the selected model, the parameter *a* was estimated individually, and then normalized by the value in non-treated condition (CON) (Fig 3A and 3B).

In the work/delay task, we used linear models to estimate the effect of remaining cost, i.e., workloads and delay, as described previously [55],

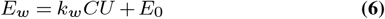

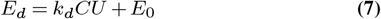

where *E_w_* and *E_d_* are the error rates, and *k_w_* and *k_d_* are cost factors for work and delay trials, respectively. *CU* is the number of remaining cost units, and *E_0_* is the intercept. We used LMMs to estimate the effect of DAR blockade on the discounting parameters. We imposed the constraint that the intercept (E0) has the same value across trials, and assumed the base statistical model in which the random effects of the two receptor types (delay and workload) affect the regression confidents i ndependently. Four models were nested to consider the presence or absence of random effects, random effects of treatment conditions, and subjects (S3 Table). The best model was selected based on BIC for the entire data set, which is the sum of the regression results for each unit faceted by individual and/or treatment condition. For example, model #1 was fit to a total of 18 data sets (3 monkeys × 3 treatment conditions (CON, MO, and HO) × 2 subtypes (D1R and D2R), and then BIC was calculated by the sum of each fitting. Modeling was performed with the lme4 package in R, and the parameters (e.g., *kw* and *kd*) were estimated from the model. We performed one-way ANOVAs to test the significance of the effect of treatment on discounting parameters with post-hoc Tukey HSD test.

LMMs were also applied for the correlation analysis between refusal rate (*E*) and reaction time (*Rt*) (Figs. 4, 6, and S5 Fig), where four statistical models were nested to take into account the presence or absence of random effects of subjects and treatment conditions, and the best-fit model was selected based on BIC (S2, S4, and S5 tables).

## Data availability

All data presented in this paper have been posted on the following public repository: https://github.com/minamimoto-lab/2021-Hori-DAR.

## ACKNOWLEDGEMENTS

We thank R. Suma, T. Okauchi, Y. Sugii, R. Yamaguchi, Y. Matsuda, and J. Kamei for their technical assistance, and K. Oyama for discussion. We also thank to Dr. MR. Zhang and his colleagues at Department of Radiopharmaceuticals Development, NIRS/QST for producing the radioligands. A Japanese monkey used in this study was provided by National Bio-Resource Project “Japanese Monkeys” of the MEXT, Japan.

