## Supporting information for "D_1_- and D_2_-like receptors differentially mediate the effects of dopaminergic transmission on cost/benefit evaluation and motivation in monkeys"

**S1 Table. Model comparison for the effect of DAR blockade on refusal rates in reward size task (for Fig 2).**

| D1R blockades (monkey KN) | | | |
|---|---|---|---|
| | model | BIC | ΔBIC |
| #1 | *E = 1/a(cond)R* | -85.6 | 9.9 |
| **#2** | ***E = 1/a(cond)R + e*** | **-95.5** | **0** |
| #3 | *E = 1/aR + e(cond)* | -93.6 | 1.9 |
| #4 | *E = 1/a(cond)R + e(cond)* | -92.6 | 2.9 |

| D1R blockades (monkey ST) | | | |
|---|---|---|---|
| | model | BIC | ΔBIC |
| **#1** | ***E = 1/a(cond)R*** | **-110.6** | **0** |
| #2 | *E = 1/a(cond)R + e* | -98 | 12.6 |
| #3 | *E = 1/aR + e(cond)* | -92.4 | 18.2 |
| #4 | *E = 1/a(cond)R + e(cond)* | -95 | 15.6 |

| D2R blockades (monkey KN) | | | |
|---|---|---|---|
| | model | BIC | ΔBIC |
| **#1** | ***E = 1/a(cond)R*** | **-126.7** | **0** |
| #2 | *E = 1/a(cond)R + e* | -120.7 | 6 |
| #3 | *E = 1/aR + e(cond)* | -103.6 | 23.1 |
| #4 | *E = 1/a(cond)R + e(cond)* | -117.4 | 9.3 |

| D2R blockades (monkey ST) | | | |
|---|---|---|---|
| | model | BIC | ΔBIC |
| **#1** | ***E = 1/a(cond)R*** | **-151.7** | **0** |
| #2 | *E = 1/a(cond)R + e* | -138.8 | 12.9 |
| #3 | *E = 1/aR + e(cond)* | -124.2 | 27.5 |
| #4 | *E = 1/a(cond)R + e(cond)* | -135.4 | 16.3 |

*a(cond)* and *e(cond)* indicate the random effects of DAR blocking treatment conditions on parameters *a* and *e*, respectively. BIC (Bayesian information criterion) is a relative measure of quality for models (#1-4). Δ BIC denotes difference from minimum BIC.

**S2 Table. Model comparison for the effect of DAR blockade on the relationship between refusal rate and reaction time in reward size task (for Fig 4).**

| Model | | D1 block | | D2 block | |
|---|---|---|---|---|---|
| | | BIC | ΔBIC | BIC | ΔBIC |
| #1 | *E ~ Rt* | **-1476.2** | **0** | **-1000.5** | **0** |
| #2 | *E ~ Rt + (Rt\|monkey)* | -1435.1 | 41.1 | -988.1 | 12.4 |
| #3 | *E ~ Rt + (Rt\|cond)* | -1443.4 | 32.8 | -964.9 | 35.6 |
| #4 | *E ~ Rt + (Rt\|monkey) + (Rt\|cond)* | -1437 | 39.2 | -980.9 | 19.6 |

*(Rt|*)* indicates random effects on regression parameters. *E*, refusal rate; *Rt*, reaction time; *cond*, treatment condition; *monkey*, subject.

**S3 Table. Model comparison for the effect of DAR blockade on refusal rates in work/delay task. (for Fig 5)**

| model | | BIC | ΔBIC |
|---|---|---|---|
| #1 | $E \sim RC + E_0 + (0 + CU\|type)$ | **4405.2** | **0** |
| #3 | $E \sim RC + E_0 + (0 + CU\|type) + (CU\|monkey)$ | 4621.5 | 216.3 |
| #2 | $E \sim RC + E_0 + (0 + CU\|type) + (CU\|cond)$ | 4745.1 | 339.8 |
| #4 | $E \sim RC + E_0 + (0 + CU\|type) + (CU\|monkey) + (CU\|cond)$ | 4858.1 | 452.9 |

*CU* and $E_0$ indicate remaining cost and intercept, respectively. *(0+ CU|*)* and *(CU|*)* indicate random effects on both regression coefficient and intercept ($E_0$) or on regression coefficient alone, respectively. *E*, refusal rate; *type*, trial type (delay or work); *cond*, treatment condition (CON, MO, and HO for D1R and D2R blocking); *monkey*, subject.

**S4 Table. Model comparison for the effect of DRA blockade on the relationship between refusal rate and reaction time in work/delay task (for Fig 6).**

D1R block

| model | | Delay | | Workload | |
|---|---|---|---|---|---|
| | | BIC | ΔBIC | BIC | ΔBIC |
| #1 | *E ~ Rt* | 1387.1 | 81.4 | **1308.2** | **0** |
| #2 | *E ~ Rt + (Rt\|monkey)* | **1305.7** | **0** | 1308.6 | 0.4 |
| #3 | *E ~ Rt + (Rt\|cond)* | 1374.1 | 68.4 | 1331.3 | 23.2 |
| #4 | *E ~ Rt + (Rt\|monkey) + (RT\|cond)* | 1329.3 | 23.5 | 1324.4 | 16.3 |

D2R block

| model | | Delay | | Workload | |
|---|---|---|---|---|---|
| | | BIC | ΔBIC | BIC | ΔBIC |
| #1 | *E ~ Rt* | 1114.8 | 38.3 | 1228.9 | 72.6 |
| #2 | *E ~ Rt + (Rt\|monkey)* | **1076.5** | **0** | 1186.2 | 30 |
| #3 | *E ~ Rt + (Rt\|cond)* | 1090.7 | 14.3 | 1191.0 | 34.7 |
| #4 | *E ~ Rt + (Rt\|monkey) + (RT\|cond)* | 1080.1 | 3.6 | **1156.3** | **0** |

*(Rt|*)* indicates random effects on regression parameters. *E*, refusal rate; *Rt*, reaction time; *cond*, treatment condition; *monkey*, subject.

**S5 Table. Model comparison for the effect of both D1R and D2R blockades on the relationship between refusal rate and reaction time in work/delay task (for S5 Fig.).**

| model | | Delay | | Workload | |
|---|---|---|---|---|---|
| | | BIC | ΔBIC | BIC | ΔBIC |
| #1 | *E ~ Rt* | **1262.2** | **0** | 1390.4 | 41.2 |
| #2 | *E ~ Rt + (Rt\|monkey)* | 1276.8 | 14.6 | 1392.4 | 43.2 |
| #3 | *E ~ Rt + (Rt\|cond)* | 1266.5 | 4.3 | **1349.2** | **0** |
| #4 | *E ~ Rt + (Rt\|monkey) + (Rt\|cond)* | 1280.1 | 17.9 | 1352.5 | 3.3 |

*(Rt|*)* indicates random effects on regression coefficient. *E*, refusal rate; *Rt*, reaction time; *cond*, treatment condition; *monkey*, subject.

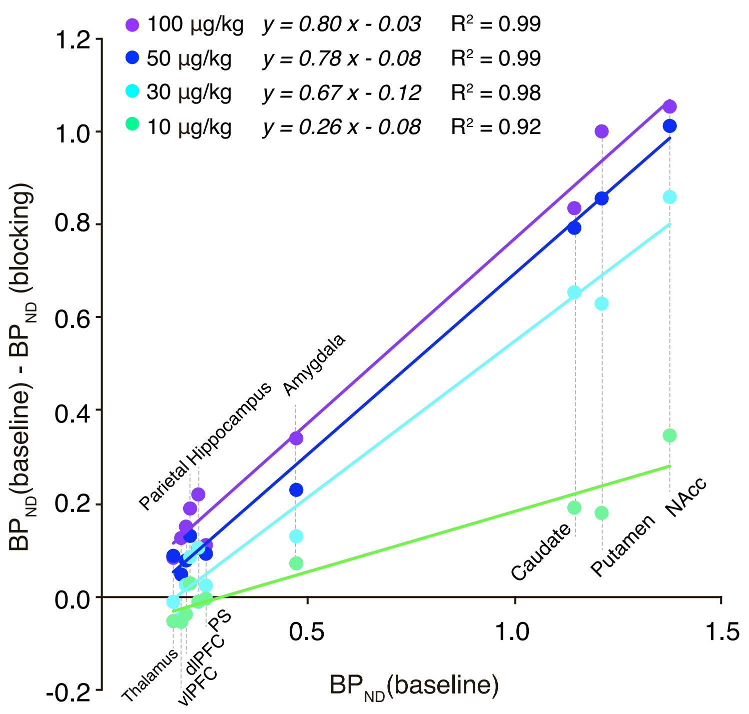


**S1 Fig. Occupancy estimation.** Example of occupancy estimation based on modified Lassen plot of [$^{11}$C]SCH23390 PET data obtained from monkey BE. Colored dots represent the relationship between decreased specific binding [i.e., $BP_{ND}$ (baseline) – $BP_{ND}$ (blocking)] and baseline [$BP_{ND}$ (baseline)] for each brain region under each blocking condition (indexed by color). Occupancy was determined as proportion of reduced specific binding to baseline, which corresponds to the slope of linear regression. In this case, $D_1$ occupancy was 80%, 78%, 67%, and 26% for 100, 50, 30 and 10 µg/kg doses, respectively. The data underlying this figure can be found on the following public repository: https://github.com/minamimoto-lab/2021-Hori-DAR.

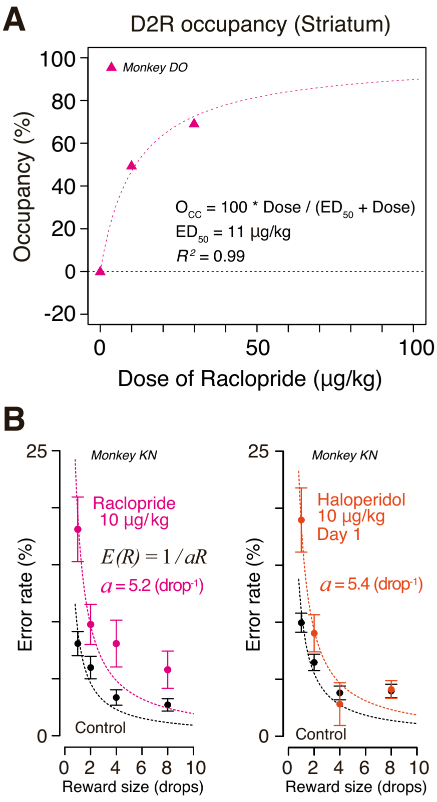


**S2 Fig. Comparable effects of $D_2R$ antagonism between raclopride and haloperidol at similar occupancy.** (A) Occupancy of $D_2R$ measured at striatal ROI is plotted against dose of raclopride. (B) Error rates as function of reward size for control (black) and after injection of raclopride (10 µg/kg, i.m. left side) and haloperidol (10 µg/kg, i.m. right side) in monkey KN are plotted. Dotted curves are best-fit inverse function (*model #1* in S1 Table). The data underlying this figure can be found on the following public repository: https://github.com/minamimoto-lab/2021-Hori-DAR.

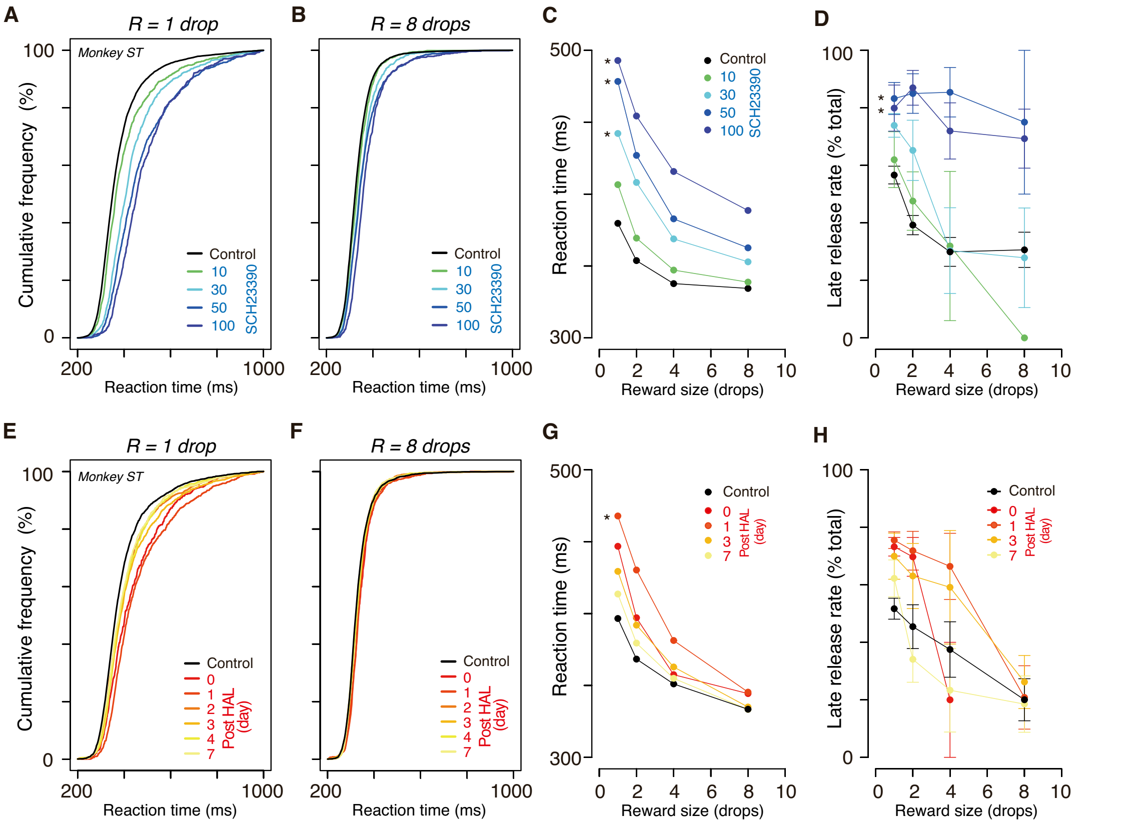


**S3 Fig. Effect of $D_1R/D_2R$ blockade on reaction time and error pattern.** (A-B) Cumulative distribution of reaction time for control and $D_1R$ blockade conditions in drop-1 and -8 trials, respectively. (C) Mean reaction time as function of reward size for control and $D_1R$ blockade conditions. Two-way ANOVA, reward x condition; main effect of condition, $F_{(4, 164)} = 109.8$, $p < 1.0 \times 10^{-15}$; main effect of reward, $F_{(3, 164)} = 111.0$, $p < 10^{-15}$; interaction, $F_{(12, 164)} = 4.7$, $p < 1.0 \times 10^{-5}$. (D) Late release rate (mean ± SEM) as function of reward size for control and $D_1R$ blockade conditions. Two-way ANOVA, reward x condition; main effect of condition, $F_{(4, 163)} = 18.6$, $p < 1.0 \times 10^{-11}$; main effect of reward, $F_{(3, 163)} = 9.8$, $p < 10^{-5}$; interaction, $F_{(12, 163)} = 1.0$, $p = 0.4$. (E-H) Same as (A-D), but for $D_2R$ blockade. Reaction time; main effect of condition, $F_{(6, 92)} = 7.2$, $p < 1.0 \times 10^{-5}$; main effect of reward, $F_{(3, 92)} = 81.9$, $p < 10^{-15}$; interaction, $F_{(18, 164)} = 0.6$, $p = 0.65$. Late release rate; main effect of condition, $F_{(6, 90)} = 3.5$, $p = 0.0038$; main effect of reward, $F_{(3, 90)} = 19.2$, $p < 10^{-9}$; interaction, $F_{(18, 90)} = 1.4$, $p = 0.14$. * significantly different from control, $p < 0.05$; post-hoc Tukey HSD. Data were obtained from monkey ST. The data underlying this figure can be found on the following public repository: https://github.com/minamimoto-lab/2021-Hori-DAR.

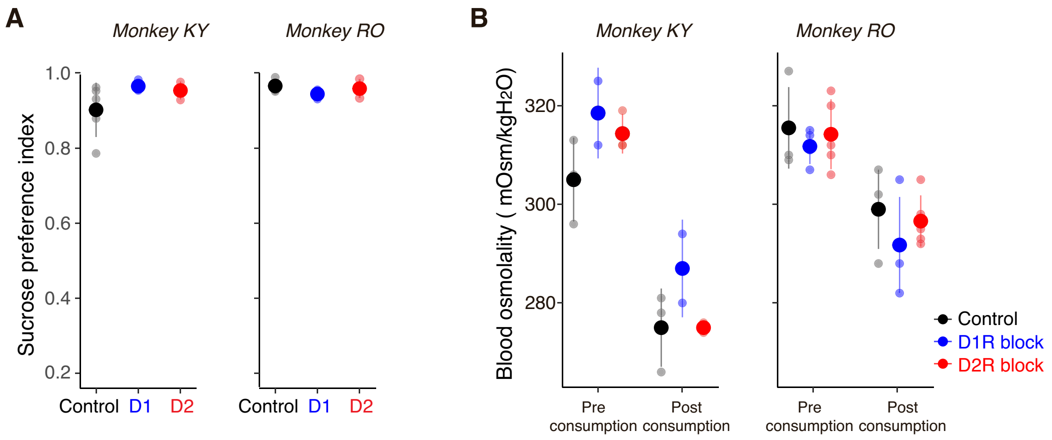


**S4 Fig. Little influence of DAR blockades on sucrose preference and blood osmolality.** (A) Sucrose preference index after administration of saline (Control), SCH23390 (30µg/kg, $D_1$) and haloperidol (10µg/kg; $D_2$, day 0), respectively. There was no significant effect of DAR blockade on overall intake (one-way ANOVA, treatment, monkey KY, $F_{(2, 8)} = 1.26$, $p = 0.33$; monkey RO, $F_{(2, 14)} = 2.01$, $p = 0.17$) or sucrose preference (one-way ANOVA; treatment, monkey KY, $F_{(2, 8)} = 1.62$, $p = 0.26$; monkey RO, $F_{(2, 8)} = 1.38$, $p = 0.31$). (B) Blood osmolality measured in serum samples obtained before (Pre) and after (Post) sucrose test. There was no significant impact of DAR blockade (2-way ANOVA, monkey KY, main effect of treatment, $F_{(2, 10)} = 4.0$, $p = 0.056$; pre-post, $F_{(1, 10)} = 93.83$, $p = 2.1 \times 10^{-6}$, interaction, $F_{(2,10)} = 0.74$, $p = 0.50$; monkey RO, treatment, $F_{(2, 20)} = 1.22$, $p = 0.32$; pre-post, $F_{(1, 20)} = 40.8$, $p = 3.1 \times 10^{-6}$, interaction, $F_{(2,20)} = 0.13$, $p = 0.88$). Filled circles and shades indicate median and raw data points, while horizontal bars indicate SD. The data underlying this figure can be found on the following public repository: https://github.com/minamimoto-lab/2021-Hori-DAR.

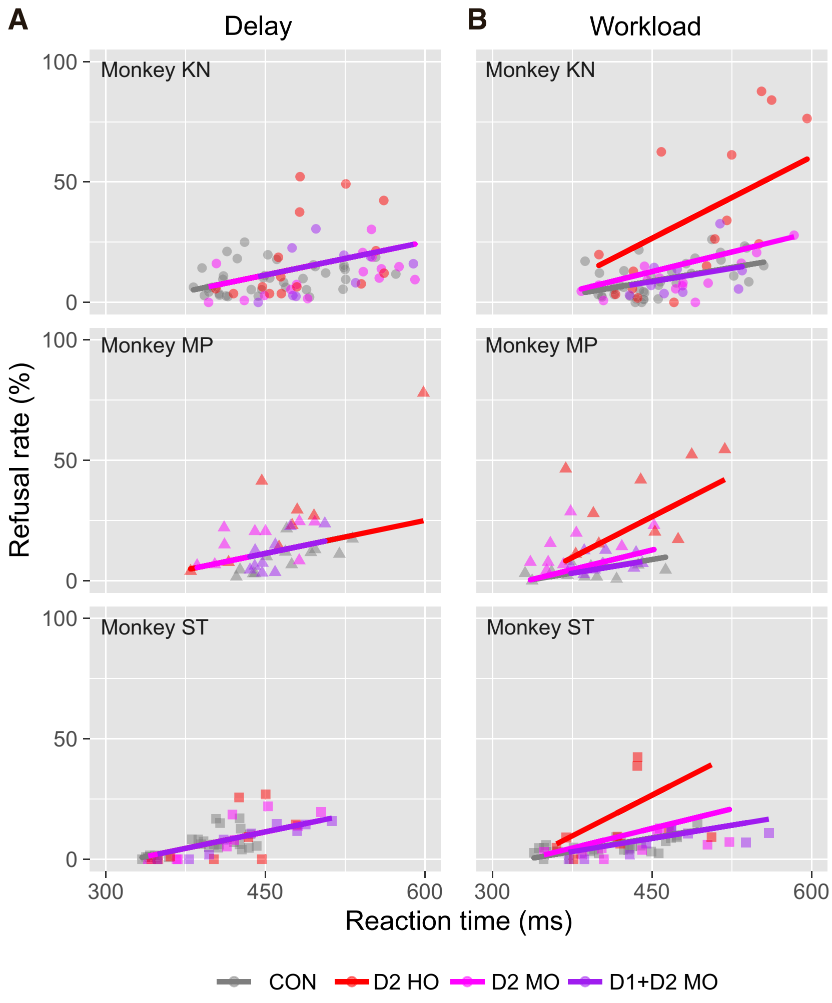


**S5 Fig. Effect of both $D_1R$ and $D_2R$ blockades on the relationship between refusal rate and reaction time.** (A) Relationship between refusal rate and average reaction time for each reward size in session-by-session for $D_2$ blocking and $D_1$+$D_2$ blocking in delay trials. Data are plotted individually for monkeys KN, MP, and ST, in order from top to bottom. Colors indicate treatment condition. Thick lines indicate linear regression lines (*model #1* in S5 Table). (B) Same as A, but for workload trials. Note that for the data in workload trials, a multiple linear model with random effect of condition (*model #3* in S5 Table) was chosen as the best model to explain the data, where the steepness of the slope under $D_1$+$D_2$ treatment was the same as that of control. The data underlying this figure can be found on the following public repository: https://github.com/minamimoto-lab/2021-Hori-DAR.
